# The human sperm head spins with a conserved direction during swimming in 3D

**DOI:** 10.1101/2022.11.28.517870

**Authors:** G. Corkidi, F. Montoya, A.L. González-Cota, P. Hernández-Herrera, N.C. Bruce, H. Bloomfield-Gadêlha, A. Darszon

## Abstract

In human sperm, head spinning is essential for sperm swimming and critical for fertilization. Measurement of head spinning has not been straightforward due to its symmetric head morphology, its translucent nature and fast 3D motion driven by its helical flagellum movement. Microscope image acquisition has been mostly restricted to 2D single focal plane images limited to head position tracing, in absence of head orientation and rotation in 3D. To date, human sperm spinning has been reported to be mono or bidirectional, and even intermittently changing direction. This variety in head spinning direction, however, appears to contradict observations of conserved helical beating of the human sperm flagellum. Here, we reconcile these observations by directly measuring the head spinning movement of freely swimming human sperm with multi-plane 4D microscopy. We show that 2D microscopy is unable to distinguish the spinning direction in human sperm. We evaluated the head spinning of 409 spermatozoa in four different conditions: in non-capacitating and capacitating solutions, for both aqueous and viscous media. All spinning spermatozoa, regardless of the experimental conditions spun counterclockwise (CCW) as seen from head-to-tail. Head spinning was suppressed in 57% of spermatozoa swimming in non-capacitating viscous media, though, interestingly, they recovered the CCW spinning after incubation in capacitating conditions within the same viscous medium. Our observations show that the spinning direction in human sperm is conserved, even when recovered from non-spin, indicating the presence of a robust and persistent helical driving mechanism powering the human sperm flagellum, thus of critical importance in future sperm motility assessments, human reproduction research and microorganism self-organised swimming.

## Introduction

Mammalian spermatozoa invariably spin as they freely swim through a fluid. Similarly to a drill, coordinated helical motion of its whip-like flagellum causes the sperm to “corkscrew” into the fluid, causing the sperm head to spin around its longitudinal axis during locomotion (David et al., 1981; Denehy et al., 1975; Ishijima et al., 1992; Linnet, 1979; Phillips et al., 1972; Rikmenspoel, 1965; Woolley, 1977). Sperm head spinning is a direct manifestation of the cyclic molecular-motor activity shaping the flagellum into a helical beat in 3D (Woolley, 1977; Woolley et al., 1984): a right-handed helical flagellum causes the sperm head to spin clockwise (CW) whilst a left-handed sperm spins in the opposite direction when seen from head-to-tail. Sperm spinning is suppressed if the flagellar waveform is purely planar (2D), as observed for human sperm penetrating viscous medium (Smith et al., 2009). The spinning ability has also been reported to be critical for a successful fertilization, related to a complex cascade of trans-membrane ion channel transport that coordinates this mode of motion (Miller et al., 2018; Zhao et al., 2022). Head spinning is thus a fundamental feature linking the molecular workings of the flagellar beat with sperm motion, and thus an important proxy of symmetry, or rather symmetry-breaking, of the helical flagellar movement in 3D (Zaferani et al., 2021).

For the past 50 years researchers have attempted to quantify and define the spinning direction of the human sperm head, though, until now, there is no consensus in the literature of its spinning directionality. Human sperm spinning has been observed to be mono-directed (Linnet, 1979; Smith et al., 2009b; Phillips, 1983; Woolley, 1977), bi-directed (Ishijima et al., 1992; Dardikman-Yoffe et al., 2020; Drake, 1974), and even intermittently directed (Bukatin et al., 2015). However, such reported variety in spinning direction appears to contradict observations of a conserved helical beating of human sperm flagellum (Bukatin et al., 2015; Ishijima et al., 1992; Linnet, 1979; Powar et al., 2022; Zhao et al., 2022), and conserved chirality of structural components in mammalian sperm flagella (Fawcett, 1975; Leung et al., 2021), with no agreement as to the direction of rotation reported in earlier studies (Woolley, 1977; Bishop, 1958; Drake, 1974; Yeung and Woolley, 1984; Woolley, 1979; Blokhuis, 1961; Daloglu et al., 2018).

The human sperm head is translucent and has an axis-symmetric morphology around its spinning axis (Figure 1), making it prone to optical illusions that may obscure accurate detection of its spinning directionality. As such, the sperm head is prone to perception bistability which can further disguise the true spinning direction in translucent objects (Liu et al., 2012). This optical illusion causes spinning translucent objects to appear to oscillate back-and-forth or spin with a switchable, and thus undefined, direction (Liu et al., 2012). At microscale this difficulty is augmented by the contrast inversion effect due to spherical aberrations in microscope objectives that cause a contrast-switch depending on the object’s relative position to the plane of focus (Keller et al., 2022; Goodman, 2005).

**Figure 1.**
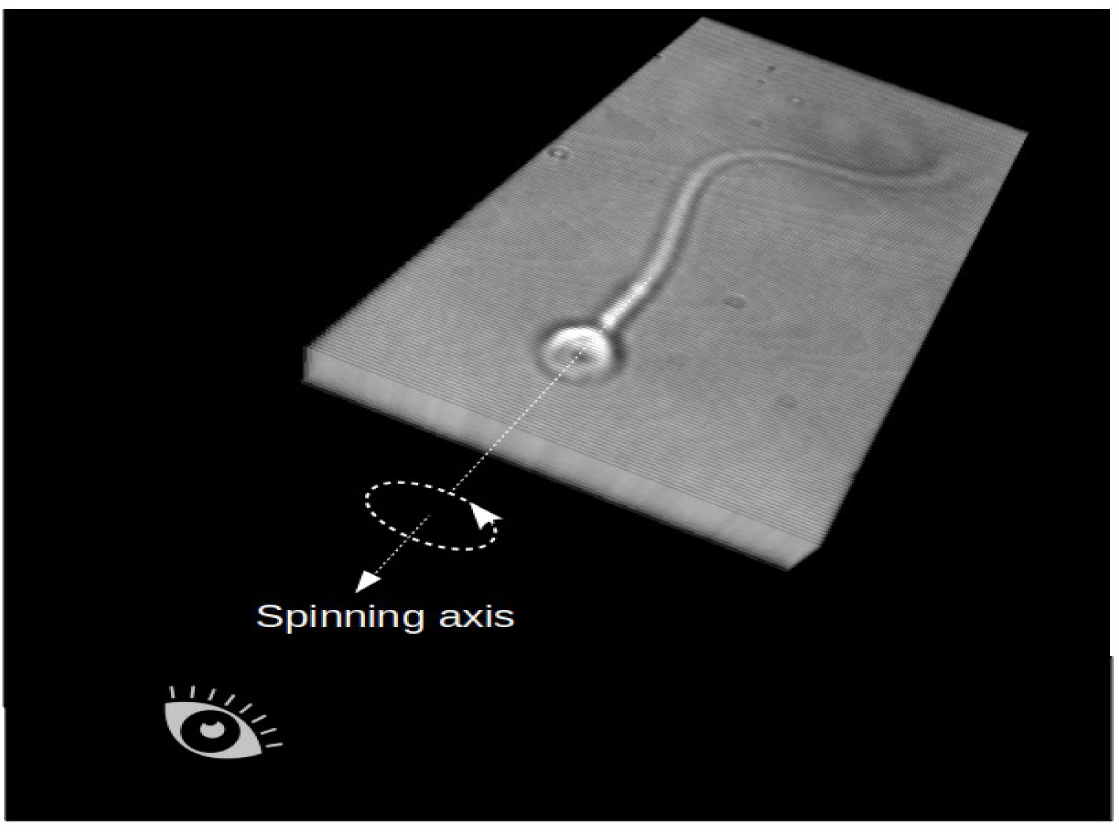
The human sperm head is translucent and has an axis-symmetric morphology around its spinning axis. The reference of observation in this work is from head to tail.

Switching the contrast of spinning objects and its subsequent dependence on the focal plane used for imaging, have unknown consequences on the detected direction of spinning (Figure 2), as we further detail in this paper. Altogether, these challenges indicate that 2D microscopy may not provide accurate measurement of the directionality of sperm head spinning (Muschol et al., 2018). Incidentally, *direct* detection of the head spinning direction in human sperm has been limited thus far to single-plane 2D microscopy. Indirect detection methods either exploit the sperm head centre-position traces (Ishijima et al., 1992) or flagelloid tracks in 3D (Ishijima et al., 1992; Bukatin et al., 2015; Dardikman-Yoffe et al., 2020), by following the trajectory of a certain point along the flagellum. In this case, the head spinning direction is assumed to follow similar rotational movement of the position traces of these counterparts. Indirect detection studies however have not yet been validated against direct measurements of head spinning, as no ground truth is available for this, highlighting this is an urgent gap in literature. Furthermore, different rotations are present during sperm swimming and it is a challenging task to infer head spinning indirectly from flagellar tracks that rotate around an average swimming axis, or from flagelloid tracers that are ill posed and cannot define robustly a common rotation point. Indeed, no modern 3D flagellar tracking research has attempted to <u>directly</u> track the head spinning direction in human sperm. This is, however, critical for understanding sperm swimming as human sperm precesses around its swimming axis: the sperm head spins at the same time that the cell as a whole rotates around its swimming direction.

**Figure 2.**
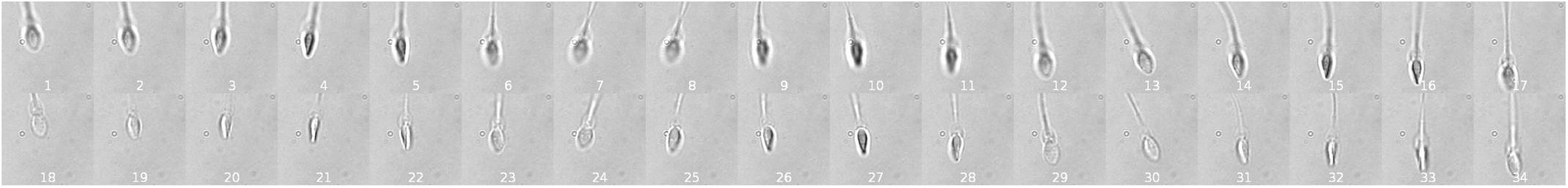
Image sequence of 17 timepoints at two different focal planes separated by 4.8 microns (image 1-17 upper focal plane, images 18-34 lower focal plane), containing the information of a complete head turn (see Video 1).

Here we solve this half-century old problem using a high-resolution multi-plane 4D method (Corkidi et al., 2008; Silva-Villalobos et al., 2014; Pimentel et al., 2012; Corkidi et al., 2021; Gadêlha et al., 2020; Hernández et al., 2022) to directly detect the head spinning direction of human spermatozoa. This method employs 4D bright-field microscopy with a 100x high-magnification objective that scans a 3D volume with high-speeds as sperm spin and swim through the fluid. The high-precision 4D microscopy described here provides a dense stack of multiple focal planes that bypass limitations of: (i) 2D microscopy that only uses a single focal plane, (ii) the image dependence on focal plane positioning, (iii) difficulties arising from contrast-switch of spherical aberrations of the lens, and (iv) perception bistability effect of translucent spinning objects. We additionally show that head sperm spinning direction cannot be confidently deduced from a single focal plane imaging. We analyzed over 400 free-swimming human spermatozoa in four different conditions, using capacitating and non-capacitating solutions within aqueous and viscous media. 100% of all sperm heads were observed to spin counterclockwise (CCW) when viewed from head-to-tail, Figure 1. Our observations show that the spinning direction in human sperm is conserved, even when head spinning is recovered from planar beating, indicating the presence of a persistent helical driving mechanism powering the human sperm flagellum. These observations may have important implications concerning the internal machinery driving the head spinning, flagellum-powered cell motility and its physiology.

## RESULTS

### a) **2D single-plane imaging cannot inform confidently the direction of head spin**

Classical 2D single-plane imaging is not suitable to establish confidently the direction of the head spin. Figure 2 shows an image sequence of 17 timepoints evolving from left to right at two different focal planes containing the information of a complete head turn. As can be appreciated, the visual information in these two planes is distinct and complex, with changes in the brightness, contrast switches and head position relative to the focal plane. Establishing the head spinning direction is not possible using single-plane information alone (see Video 1). Visual inspection by different observers leads to different spinning directions due to bistability perception.

**Video 1.**
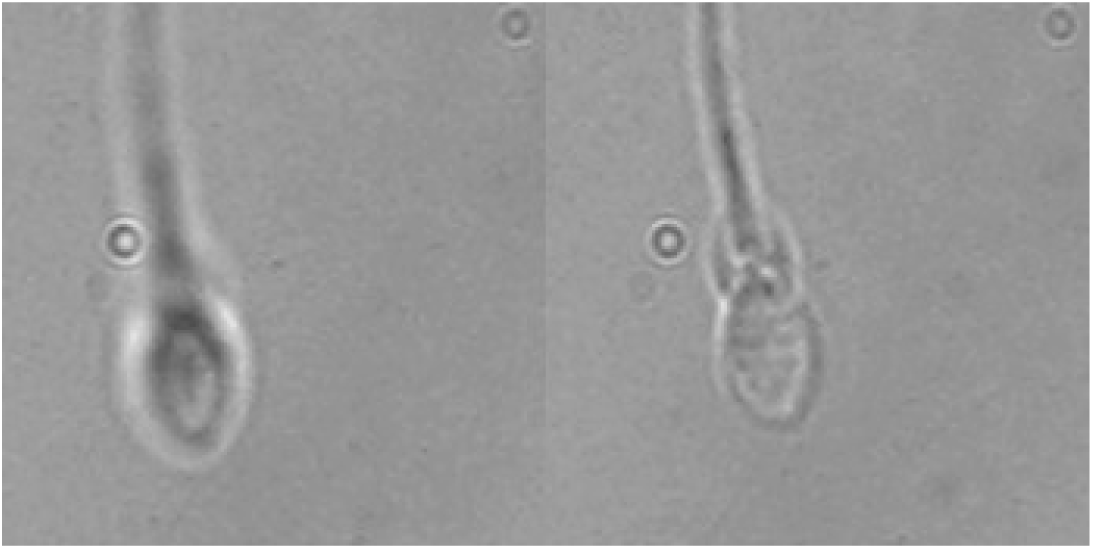
Establishing the head spinning direction is not possible using single-plane information alone. Visual inspection by different observers leads to different spinning directions due to bistability perception.

### **b)** 3D bright field image stacks acquisition and contrast inversion for the human sperm head

Figure 3 shows 25 consecutive focal planes (out of 50) from a single piezo rising slope and assumed to correspond to a single instant. The experimental setup provides a temporal resolution of 1/160 seconds and a spatial resolution along the *z* direction of 0.4 mm. In this figure, different head elements and their contrast-inversion are revealed as the focal plane moves through the sperm head, appearing as bright pixels inside the head with a dark halo outside to the head behind the focal plane (Figure 3(1)), and as dark pixels inside the head with a bright halo outside to the head in front of the focal plane (Figure 3(25)). Note particularly how the left narrowest sides of the sperm head begin to appear as a bright white border (as detailed in the next sub-sections) when images move farther from the objective lens (15 to 11) in Figure 3. This contrast inversion effect will be explored in the following section to track head spin directionality.

**Figure 3.**
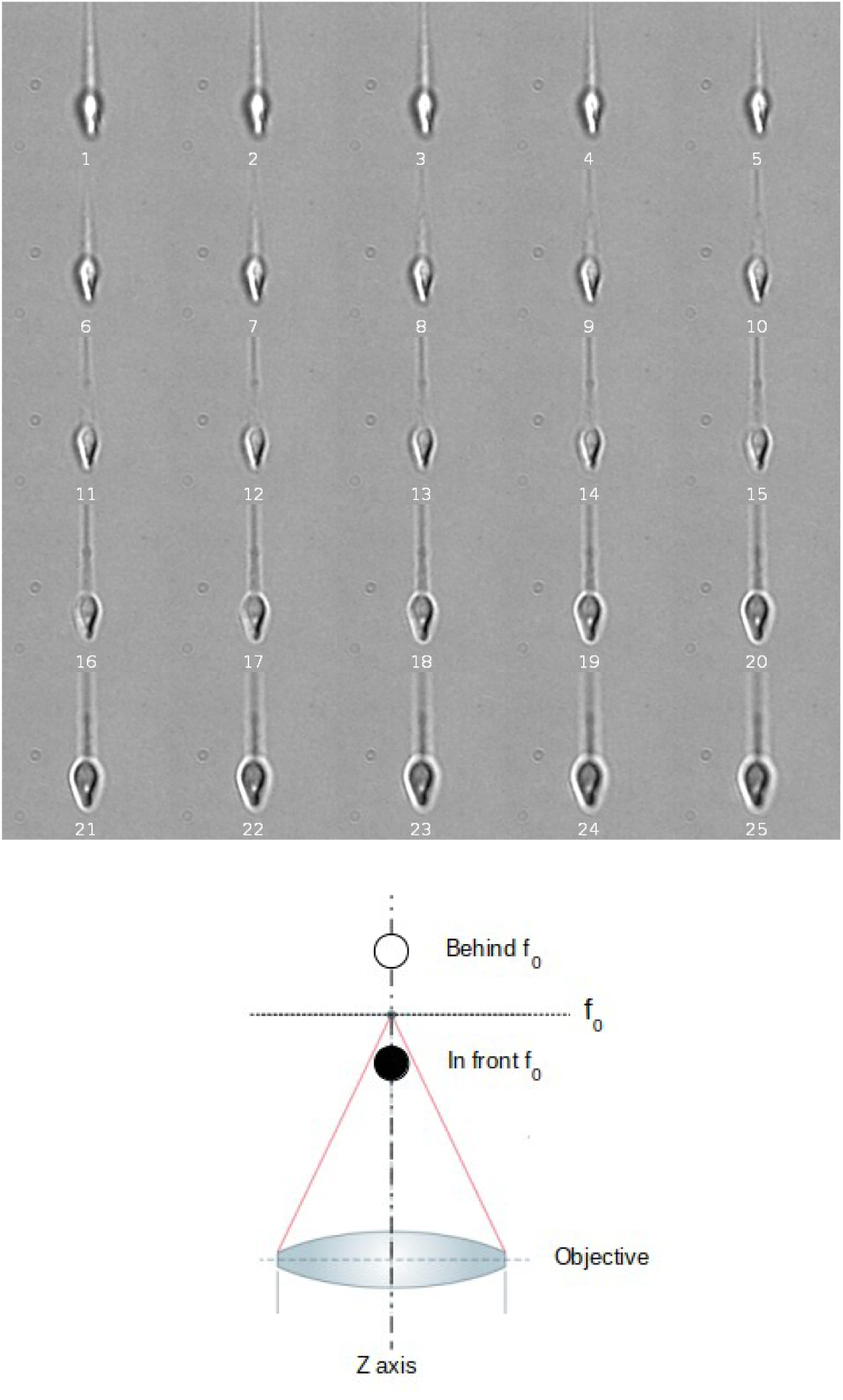
Multifocal plane acquisition. From left to right and top to bottom, this figure shows 25 consecutive focal planes (out of 50) acquired while the piezoelectric device is rising. Frames (150 x 150 pixels each) were cropped, for visualization purposes, from 640 x 480 pixels image sequences. Each focal plane is separated by 0.4 mm respectively (see Materials and Methods). The schematic at the bottom depicts how the spherical aberration of the optical system produces a contrast inversion over translucent objects depending on their position with respect to the focal plane f_0_. The object appears dark if placed in front of the focal plane f_0_, while bright if placed behind.

### **c)** Contrast inversion of the sperm head is due to the spherical aberration of the objective lens using bright-field microscopy

The Rayleigh-Sommerfeld back propagation reconstruction method has been commonly used to reconstruct the sperm flagellum in 3D (Bukatin et al., 2016), but this is based solely on diffraction and cannot distinguish between defocus before or after the focused object plane (Lee et al., 2007). Usually, the contrast inversion of the defocused object is used heuristically to post process the results of the back-propagation method to produce a final reconstructed object (Lee et al., 2007), alternatively, phase information from the reconstructed object can be used to post-process the reconstruction itself (Wilson et al., 2012). It is well known that microscope objectives are extremely well corrected for conjugate planes, which are focused objects and image planes, but this is not true for non-conjugate planes. Aberrations, particularly spherical aberrations, can strongly contribute to image quality for non-conjugate planes (Keller et al., 2022; Kidger, 2002). Here we demonstrate that it is the spherical aberration of the objective lens that produces a contrast inversion as a function of the position of the object relative to the focused object plane (Figure 3).

Diffraction effects that appear in the point spread function (PSF) are given by the Fourier transform of the lens aperture function multiplied by the aberration function,

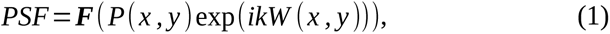

where ***F***() is the Fourier transform in the lens aperture plane, (*x, y*) is the position in the image plane, *P*(*x, y*) is the lens pupil (almost always taken as a circle), *k* =2 *π* /*λ* is the light wave number, and *W* (*x, y*) is the wave aberration function given by (Goodman, 2005)

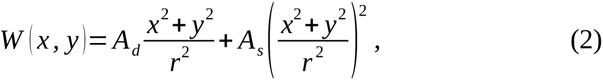

where *A_d_* is the amplitude of the defocus, *A_s_* is the amplitude of the spherical aberration, and *r* is the radius of the lens pupil. The first term in Equation 2 is the defocus contribution, and the second term is the contribution of the spherical aberration. One important aspect is the sign of the defocus term. From Goodman 2005, using the Gauss equation for a defocused system we have:

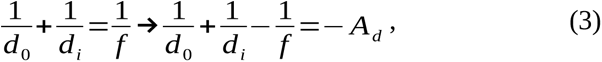

so that *A_d_* =0 when the system is focused. Given that *f* and *d_i_* (focal length and the distance from the lens to the image) are fixed, and the distance from the object to the lens (*d_0_*) changes when the optical system is defocused, when *d*_0_ increases (the object is further away from the objective lens), the term 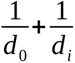 is smaller and *A_d_* > 0 . When *d*_0_ decreases (the object is closer to the objective lens), the term 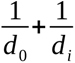 increases and *A_d_<*0. We assume that the spherical aberration has the same sign for a defocus below and above the geometrical focal plane. Simulations for a rectangular object are shown in Figure 4. The diffraction from defocus alone in Figure 4A and B, does not give an inversion of contrast between the object positions below and above the focused object plane. Figure 4C and D, however, where a small spherical aberration contribution has been included, does show the contrast inversion effect. Finally, Figure 4E shows the case of an inclined object where the left part of the object is further away from the objective lens and the right part is closer to the objective lens, showing the contrast inversion within the object. It is important to note that a change in sign of the spherical aberration contribution changes the contrast inversion: with *A_s_* > 0 an object closer to the objective lens than the focused object plane would be dark and an object further from the objective lens would be bright. The sign of the aberration depends on the specific design of the optical system and so could vary between different laboratories working with different objectives. Our simulations consider a spherical aberration within the range of the optical system used in our experiments.

**Figure 4.**
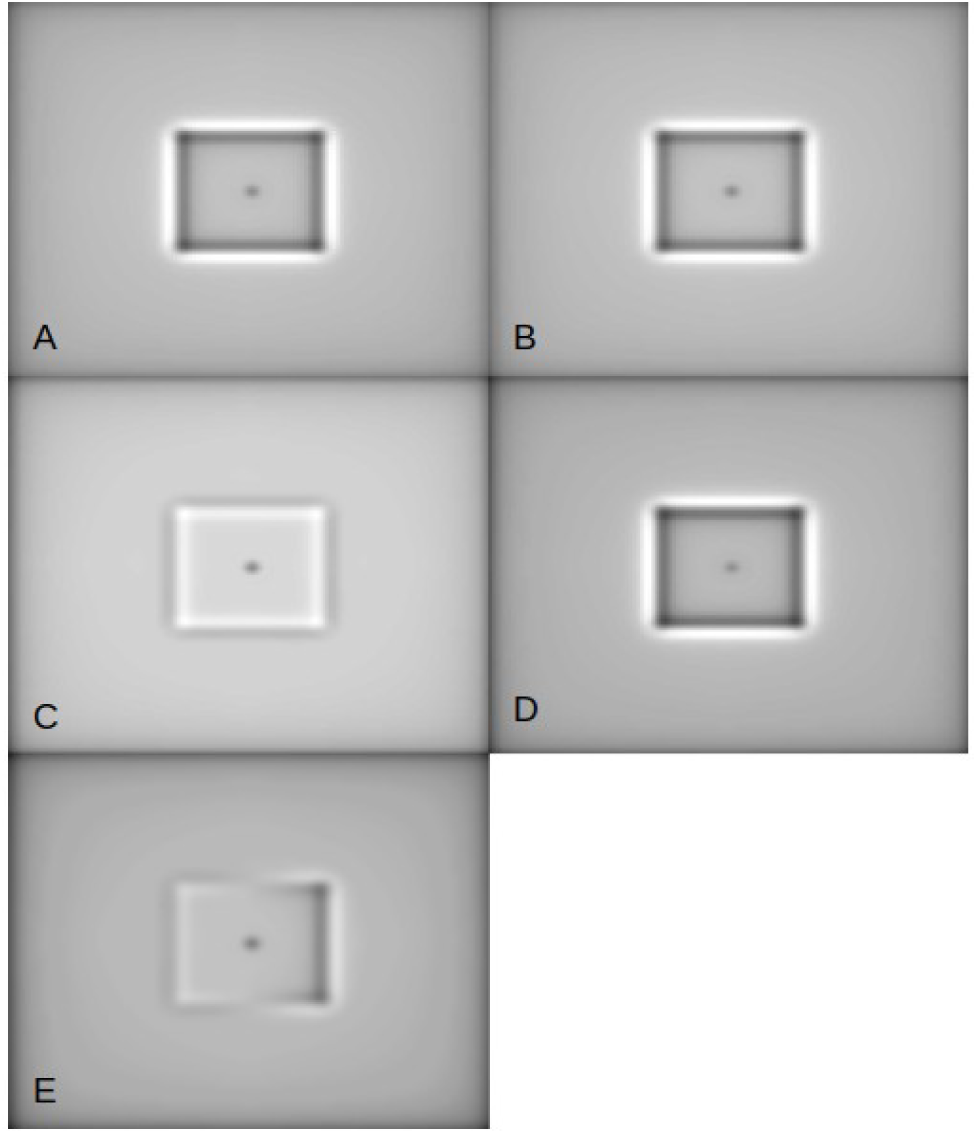
Results of the simulation of image formation for a weakly scattering phase object. (A) image with *A_s_*=0 and *A_d_* =0.3 ; (B) image with *A_s_* =0 and *A_d_* =−0.3; (C) image with *A_s_*=−0.0004 and *A_d_* =0.04; (D) *A_s_* =−0.0004 and *A_d_* =−0.04 ; (E) image for an inclined planar object with *A_s_*=−0.0004. The centre of the object plate is in the focused object plane of the lens and the right part of the object is closer to the objective lens.

The contrast-switch inside the object is a direct manifestation of the object’s inclination relative to the objective lens (Figure 4E). Here we exploit for the first time this unique optical effect to extract the directionality of a spinning object, such as the sperm head, as detailed below. Figure 5 shows how this contrast-switch of its planar projection varies when the inclination of the object increases as the object spins around its long axis. Clockwise spinning causes the bright part (further away from the objective lens, see white arrows) of the inclined object to always move from right-to-left and the dark part to move from left-to-right (dark arrows), whilst counterclockwise spinning causes the bright region to always move from left-to-right and at the same time that the dark region moves from right-to-left. Once a half-cycle is completed, and the flat object is parallel to the objective lens, the bottom part of the object reaches the top and vice versa, re-setting the contrast inversion: any part of the object reaching the top appears as bright, likewise any part of the object reaching the bottom will appear as dark. In other words, after a half-cycle, a bright/dark region abruptly re-appears at starting location, after reaching the end of the object on the opposite side. For this reason, if an object spins in the same direction, this will be manifested in the image as the bright or dark regions of the objects moving with a persistent direction relative to the object’s orientation (left-to-right or right-to-left). Likewise, if the object spinning direction is reversed, the direction of motion of bright/dark regions will equally reverse relative to the object’s orientation (right-to-left or left-to-right). The spherical aberration effect causing the contrast switch within the object’s image allows for robust, and yet simple, detection of spinning direction of axis-symmetric objects, such as the human sperm head demonstrated below, that otherwise would not be possible, and this feature is exploited here for the first time.

**Figure 5.**
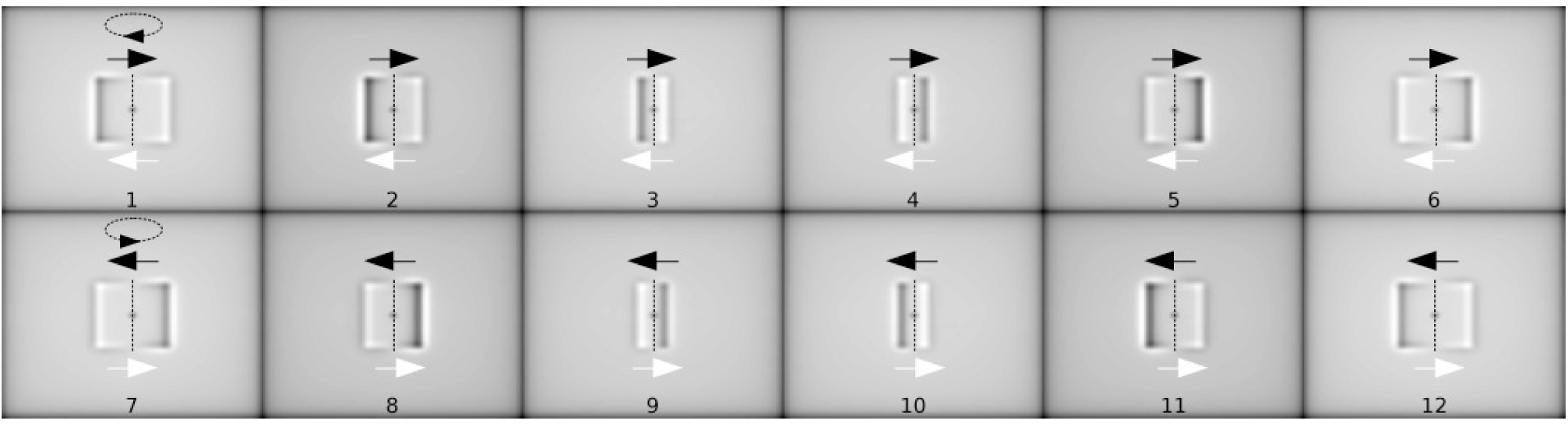
Simulation results of a spinning object, showing the direction of movement of the dark and bright left/right edges from 2D imaging from a fixed focal plane projection. Top: (images 1 to 6) a clockwise rotation viewed from the top of the images; bottom: (images 7 to 12) a counterclockwise rotation. A_s_=-0.004 and the maximum defocus is A_d_=*±*0.2.

### **d)** Bright region induced by sperm head inclination moves with head spinning

The human sperm head has a flattened side, and when the head spins around its longitudinal axis, the narrow side behind the focal plane is clearly seen as a bright region, whilst the part of the head above the focal plane appears as dark, when the sperm head is inclined relative to the objective lens (Figure 6, images 4-6, see arrows). The contrast switch inside the sperm head is due to the head inclination relative to the objective lens (images 4-6); when the sperm head is parallel to the objective lens, no switch in contrast occurs (images 8-9), as demonstrated in the above section. The time-sequence in Figure 6 (images 4-7), shows the bright region moving from left-to-right relative to the sperm head orientation due to changes in inclination of the sperm head caused by the head spinning.

**Figure 6.**
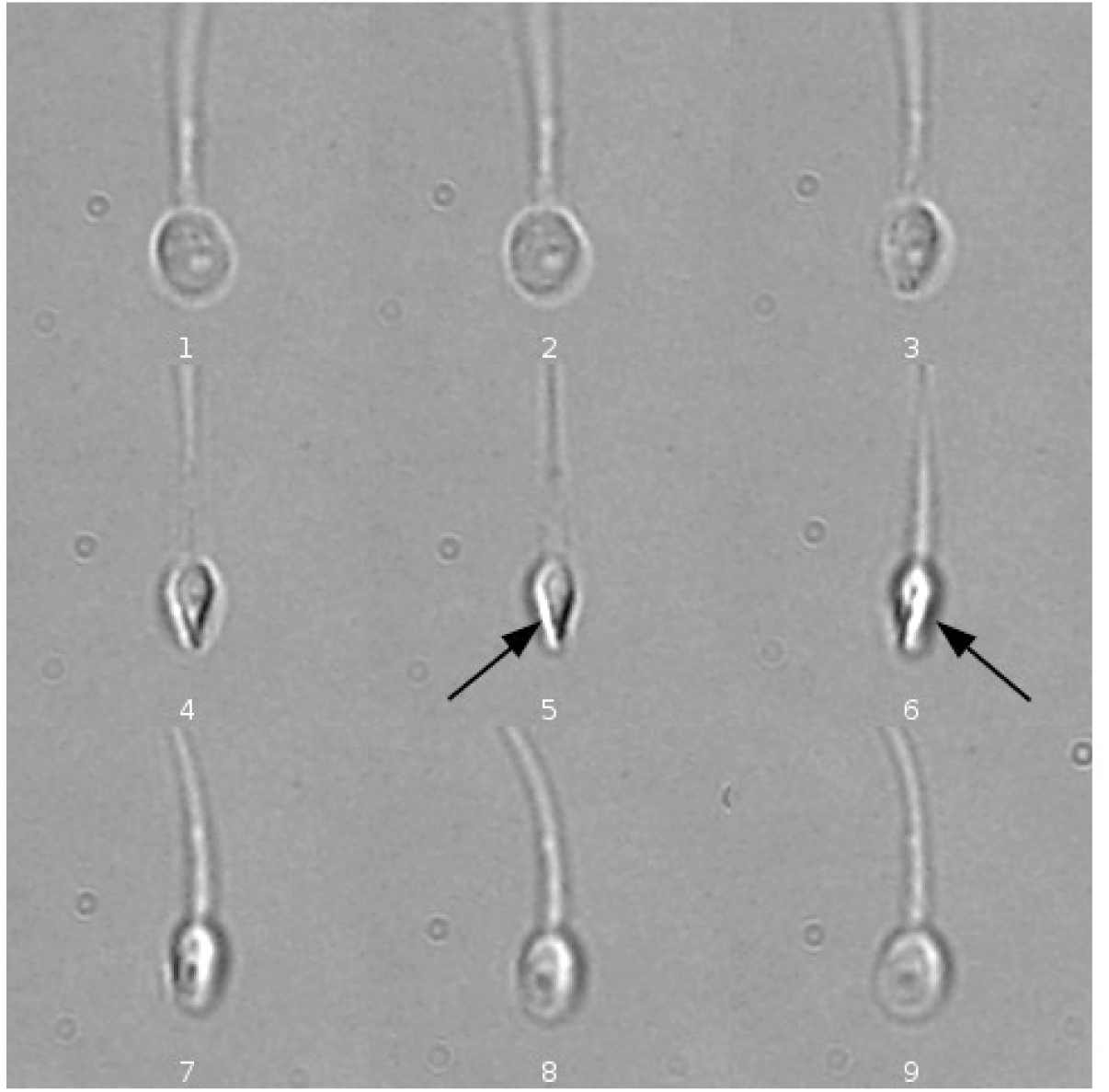
The contrast inversion optical effect during a half turn of a sperm head. Nine consecutive timepoints where the contrast inversion optical effect is clearly observed (same focal plane), i.e. the narrowest, focally lower, border of the sperm head (behind the focal plane) is enhanced resulting in a bright border while the sperm head is inclined during head spinning (highlighted with arrows).

This switch in contrast when the head is inclined is barely perceptible when observed with 2D microscopy (Figure 2), as the sperm head continuously changes its position relative to the focal plane when swimming freely in the fluid (see Video 1). A minimal focal change at the micron scale is sufficient to prevent the detection of bright-dark regions induced by the sperm head inclination, as shown in Figure 4E. Indeed, Figures 2 and 6 show the inversion in contrast due to inclination of the sperm head is clearly visible for only a few focal planes as the piezo rises. It would be a challenging task to change a single focal plane dynamically, at the microscale, to keep the sperm head exactly in focus to enable the detection of this change in contrast optical effect across the head using 2D microscopy, as shown in Figure 5. Our multifocal system bypasses this challenge and allows unique detection of the accumulated changes in contrast from bright-to-dark regions induced by the sperm head inclination from a stack of multiple focal planes (Figure 3), as detailed below.

### **e)** Validation of the sperm head spinning detection using contrast switch of the sperm head

We validated the use of the contrast inversion of the sperm head shown previously to detect the head spinning direction against direct tracking of the motion of a particle attached to a sperm neck during head spinning. Figure 7 shows a sperm cell with such a particle rigidly attached to its neck, whilst spinning 360 degrees during free-swimming motion. In the time-sequence for the Figure 7A, the focal plane is approximately placed between the sperm and the particle along *z*, in such a way that when this particle is behind the focal plane (images 1-4), it appears as bright, while when the particle is in front of the focal plane (images 5-8), it appears as dark. The arrow on the particle shows the direction of the displacement during head spinning; the black arrow indicates that the particle is behind the focal plane (bright particle), while the red arrow indicates when the particle is in front of the focal plane (dark particle). As such, the particle rotates in the CCW direction following the sperm head spinning motion: the particle moves from left-to-right when behind the focal plane (1-4), and moves from right-to-left when above the focal plane (5-8). On the other hand, the bright border arising from the contrast inversion when the sperm head is inclined relative to the objective lens always moves in the same direction relative to the sperm head orientation, from left-to-right, as shown by the arrow on this bright region, regardless of whether the particle is above (5-8) or below (1-4) the focal plane in each of its half-cycle (see Video 2). CCW spinning (as seen from head-to-tail) causes the sperm head bright region to always move from left-to-right relative to the sperm head, in agreement with the direction of the particle rotation above.

**Figure 7.**
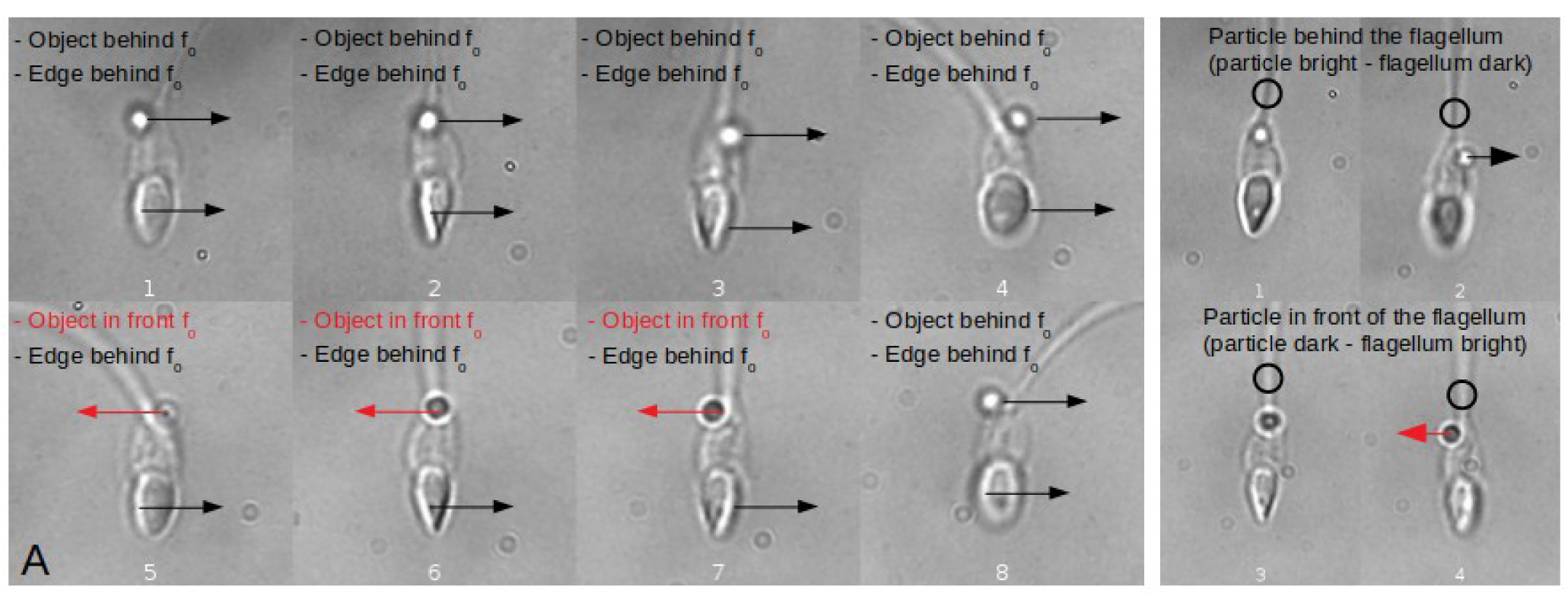
Tracking a translucent particle stuck on a spermatozoon’s neck, presenting the contrast inversion optical effect while turning 360 degrees with a free-swimming sperm. (A) Focal plane between Particle and Bright border: when the particle is above the spermatozoon’s neck it appears bright (behind the focal plane), while appearing dark when under the neck (in front of the focal plane); simultaneously, the border of the spermatozoon is always behind the focal plane appearing bright, showing a CCW spin direction when seen from head-to-tail. Note the change of direction of the particle when it is dark, as the particle rigidly follows the head spinning. (B) Focal plane between Particle and Flagellum: when the focal plane is placed between the particle and the flagellum, it becomes evident their relative position: in the sequence 1-2 (top images), the particle (bright) is located behind the flagellum (dark) while moving to the right, while in the sequence 3-4, the particle is in front (while moving to the left). A CCW spin is evident from these images. This shows that the proposed method is capable of detecting spinning directions.

When the focal plane is placed between the particle and the flagellum (Figure 7B), it becomes evident their relative position: in the sequence 1-2 (top images), the particle (bright) is located behind the flagellum (dark) while moving to the right, while in the sequence 3-4, the particle is in front (while moving to the left). The movement direction of the particle, combined with its relative position to the flagellum makes to confidently conclude a CCW spin.

**Video 2.**
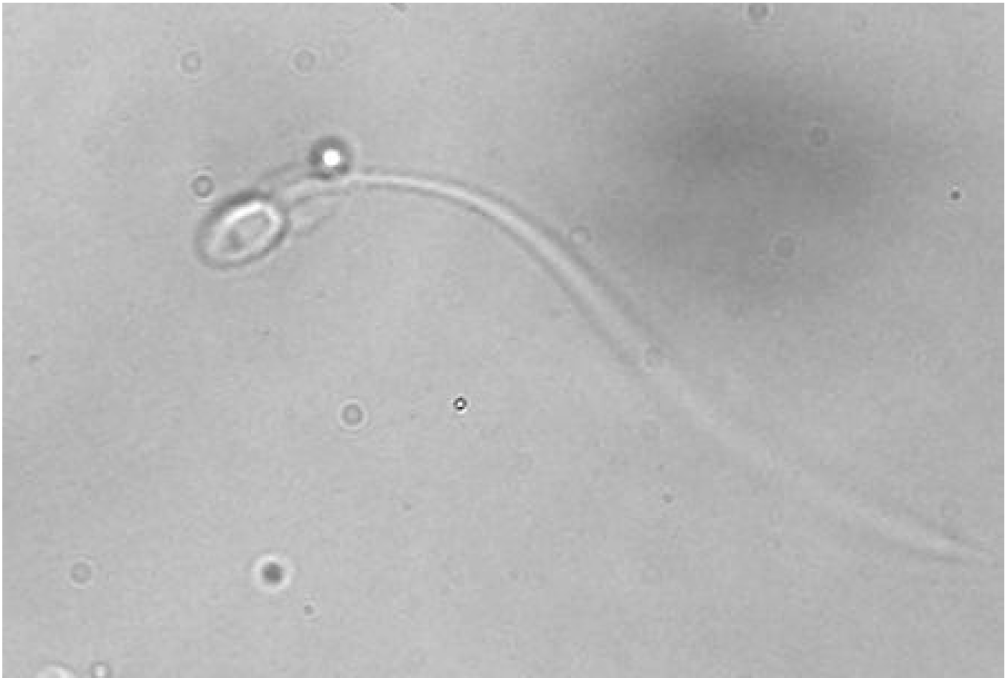
A sperm cell with a translucid particle rigidly attached to its neck, whilst spinning 360 degrees during free-swimming motion.

### **f)** Detection of sperm head spin direction from multifocal stacks

Here, we exploit the volume stack and accumulate the contrast-switch caused by the head inclination occurring at different focal planes by integrating this multifocal plane information in a single image. This circumvents the fact that the contrast switch across the sperm head is only observable for a few focal planes, as shown above. To this purpose, we calculated a 2D Maximum Intensity Projection (MIP) image (Schindelin et al., 2012) for each piezo rising slope *z* stack containing 50 focal planes through the whole acquisition time of 3.5 sec. The 2D MIP of the z-stack accumulates in a single image the maximum values of all the focal planes, i.e. bright regions in the image shown in Figure 6 for a time sequence.

The integrated bright region induced by the switch in contrast when the sperm head is inclined is manifested as a superimposed one-sided halo, only present when the sperm head is inclined relative to the objective lens (images 4-7). Figure 8 shows the accumulated bright region moving from left-to-right relative to the head as time progresses from time sequence 4-7 (see also Video 3). The motion of the accumulated MIP bright feature over the course of time follows the direction of head spinning, as demonstrated in previous sections.

**Figure 8.**
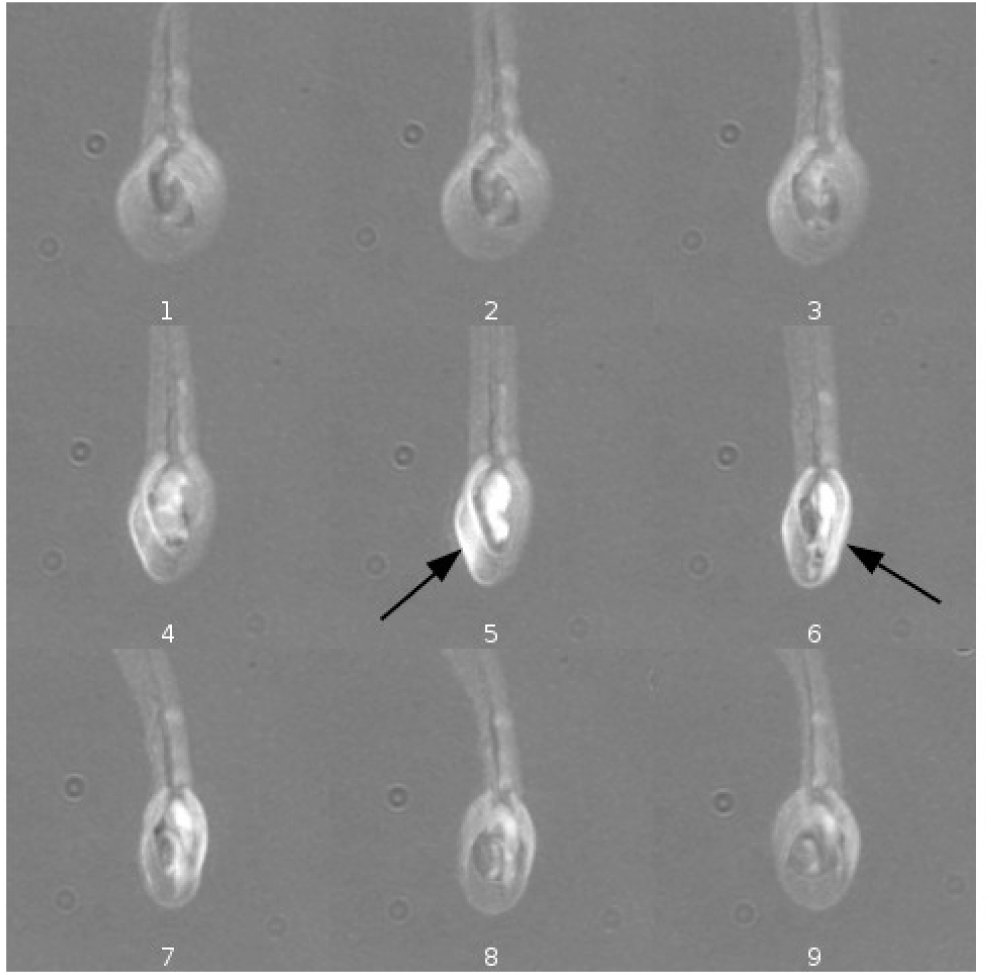
2D MIP’s (Maximum Intensity Projections) of 9 consecutive piezo rising slopes - timepoints- (50 images each projection, images 1 to 9) during a half turn of a sperm head (left to right, top to bottom). The brightest pixels over the z axis from 50 images corresponding to one half cycle of the piezo device are projected into a single plane to integrate the spherical aberration spread over the z range. The bright region moves from left-to-right relative to the sperm head (arrows) (See Video 3).

**Video 3.**
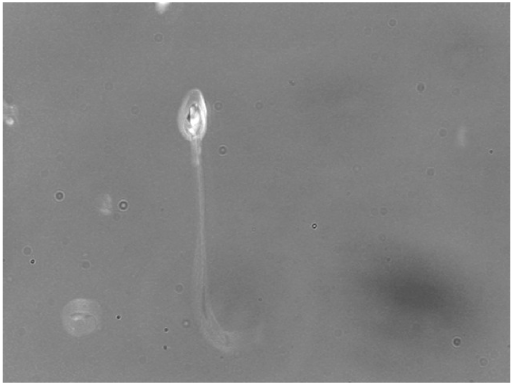
The 2D MIP of the z-stack accumulates in a single image the maximum values of all the focal planes, i.e. bright regions for a time sequence. A bright region moves from right-to-left relative to the head as time progresses. At the final part of the video, another spermatozoon appears swimming in the opposite direction, showing the bright region to move consistently in the inverse direction (see also Supporting Information and Figure S1).

The directional motion of the superimposed bright regions is extracted from the variations on the intensity profile moving along the segment ***bb’*** (red line shown in Figure 9). To detect the direction of the movement in which the one-side bright halo moves with respect to the longitudinal axis of the head (as placed in front of the head -viewing the sperm from the tip of the head to the flagellum-, line **C** Figure 9A), we have employed the following steps:

**Figure 9.**
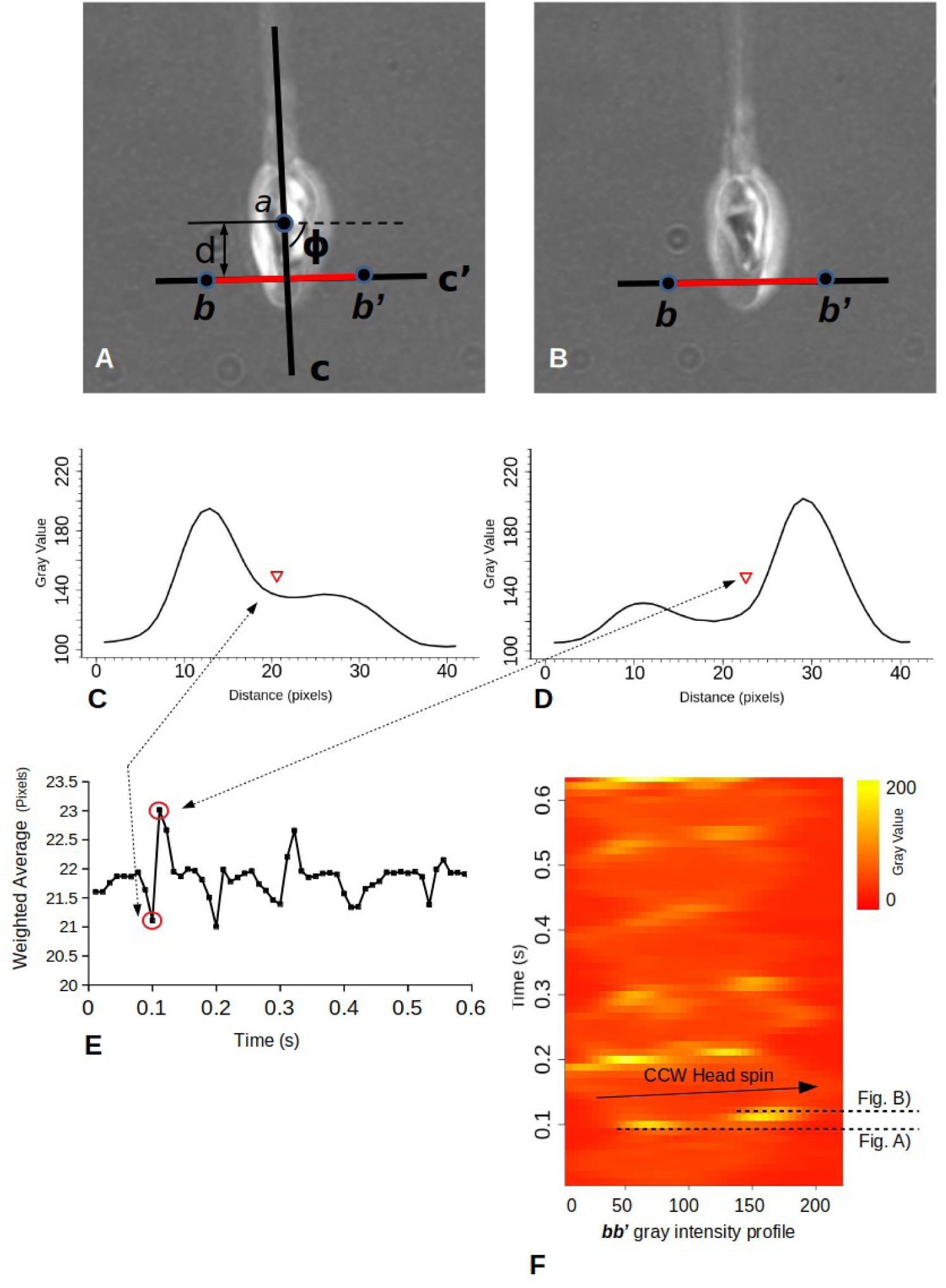
Sperm head bright region travels with a conserved direction. (A-B) The tracking process estimates the angle ϕ of the dominant direction of the spermatozoon relative to the microscope frame of reference and defines the orientation of the sperm (line **C**, see Corkidi et al., 2021). The straight-line **C’** is perpendicular to **C**, and the intersection point is at a distance of d mm from head center (1/3 the average size of the long axis of the human sperm head). ***b*** and ***b’*** are located symmetrically along **C’** at a distance ***b*** from the intersection of **CC’**. The coordinates of ***bb’*** (represented by the red line) are used to measure the intensity profile over the 2D MIP image (see Video 4). (C-D) The corresponding gray level of the 2D MIP profiles (from A-B) with its center of mass depicted by the red triangle. (E) Weighted average of 2D MIP profiles for two and a half head turns. Minima denote that the weighted average of the intensity profile is shifted towards the ***b*** side of the sperm head (taking ***b*** (from ***bb’***) as the origin for the profile) while maxima denote the shift towards the opposite ***b’*** side. The transition from a minimum to a maximum corresponds to half a turn of the sperm head in a CCW direction (two and a half CCW turns for this series). (F) Intensity profiles (***bb’***) kimograph; dashed lines point out the bright profiles corresponding to Figures A and B denoting a 180 degrees head turn. The lower time ascending arrow show waves of the brightness level traveling from left-to-right, indicating CCW head spin when seen from head to tail.

1) To track the sperm head position, we used the method described in (Corkidi et al., 2021). This method provides the spermatozoa dominant orientation over time (angle ϕ in Figure 9A) relative to the microscope fixed frame of reference, depicted by the line **C**. The center of the sperm head over line **C** is defined as *a*. These two experimental parameters define the position and orientation of the major axis of the sperm head represented by **C** (see Corkidi et al., 2021). By construction, **C’** is perpendicular to **C**, and the intersection point is at a distance of d mm from *a* (1/3 the average size of the long axis of the human sperm head). Finally, ***b*** and ***b’*** are located symmetrically along **C’** at a distance ***b*** from the intersection of **CC’**. The pixels along ***bb’*** (represented by the red line) are used to measure the intensity profile over the 2D MIP image (see Video 4). The direction of motion of the maximum intensity of the profile series (by using variation in position of the weighted average -see next section-along the segment bb’, as shown in Figure 9C and D) defines the sperm head spin direction. The equation describing the points along the line **C** is *y*=(*tanϕϕ*)*x*+(*y_a_*−*x_a_ tanϕϕ*), the intersection point between **C** and **C’** is given by the condition 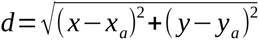 and its coordinates (*x_d_, y_d_*). The equation describing the points along **C’** is defined by 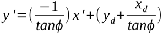. The points ***b*** and ***b’*** are positioned at distances 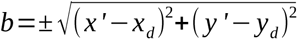 along **C’** centered at the intersection with **C**. The set of points that are embedded in the ***bb’*** segment are 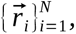, where *N* is the number of points belonging to ***bb’*** (red line in Fig 9.). From this, the gray level profile over the segment ***bb’*** can be quantified.

## 2) Determination of the sperm head spinning direction

The position of weighted average of each intensity profile along ***bb’,*** while the head is turning, is quantified to establish the head spin direction. The weighted average position over ***bb’*** is 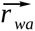, analogous to the “center of mass” of the intensity levels along ***bb’***. Tracking 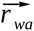 over time reveals the movement of the brightest part of the sperm head 2D MIP. This feature applied to the gray level profile over the line segment ***bb’*** (see Figure 9) is expressed as: 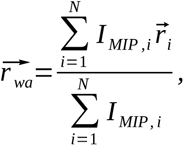, where *I _MIP,i_* is the gray level value of the corresponding 2D MIP image ⃗*r_i_*. If ⃗*r_wa_* moves from ***b*** to ***b’*** the cell is rotating CCW and otherwise CW, as shown in Figures 5 and 9.

Figure 9A and B shows the segment ***bb’*** positioned over the sperm head in two consecutive time points with the bright border moving from ***b*** to ***b’***. The corresponding gray level of the 2D MIP profile is plotted at the bottom of each image in C-D. Figure 9E shows the values of the weighted average for two and a half head turns as shown in Video 3 and Video 4. Red circles over the first minimum and maximum correspond to the first half turn. Minima denote that the weighted average of the intensity profile is shifted towards the ***b*** side of the sperm head (taking ***b*** (from ***bb’***) as the origin for the profile), while maxima denote the shift towards the opposite ***b’*** side. Figure 9F consists of the intensity profiles (***bb’***) kimograph; lower dashed lines point out the bright profiles corresponding to Figures A,B,C,D denoting a 180 degrees head turn. The upper time ascending arrow from left-to-right of the kimograph denotes a CCW head turn.

**Video 4.**
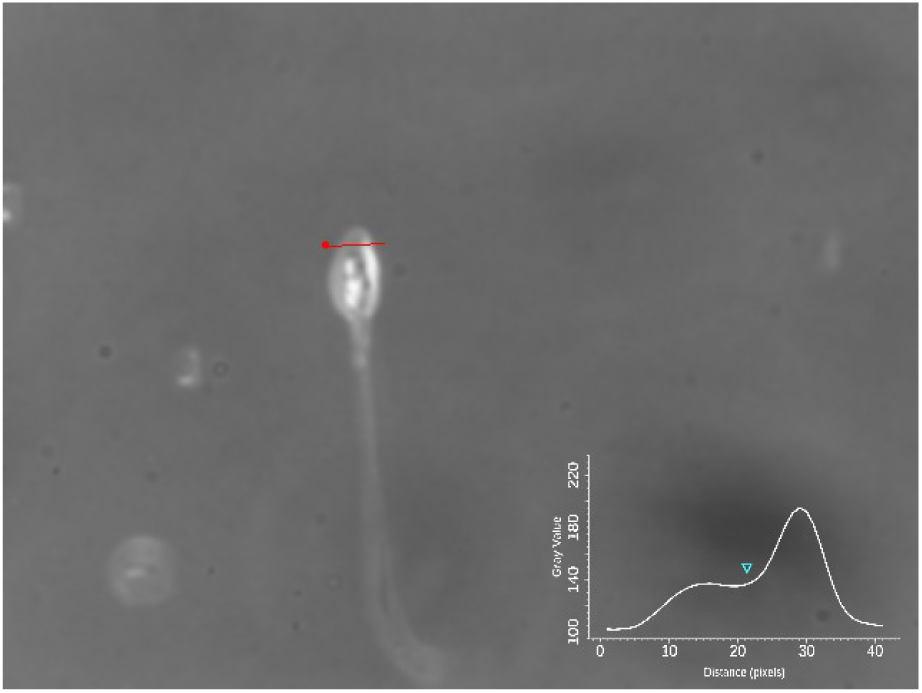
Gray level of the 2D MIP profiles (over the tracked red line ***bb’*** from Figure 9) with its center of mass depicted in the graph by the blue triangle.

### *g*) Human sperm spin with a conserved counterclockwise direction

We acquired 409 human spermatozoa while freely swimming in 3D; 180 under non-capacitating conditions and 229 in capacitating conditions. Both groups were studied in aqueous and viscous media (see Materials and Methods). We found that in aqueous media, regardless of the medium used (non-capacitating or capacitating), all the sperm spun in the CCW when seen from head to tail and swimming freely. In contrast, in viscous and non-capacitating medium, 57% of the analyzed sperm were not spinning at all, while the remaining 43% spun in the CCW direction. Interestingly, in capacitating conditions these percentages inverted: 22% of the analyzed sperm did not spin at all, whilst 78% spun with the conserved CCW direction. The results summarized in Table 1 suggest that capacitating media also influences the ability of sperm to spin in a viscous fluid. All human sperm observed to spin did so by turning in the CCW direction when seen from head to tail regardless of the experimental condition.

**Table 1.**
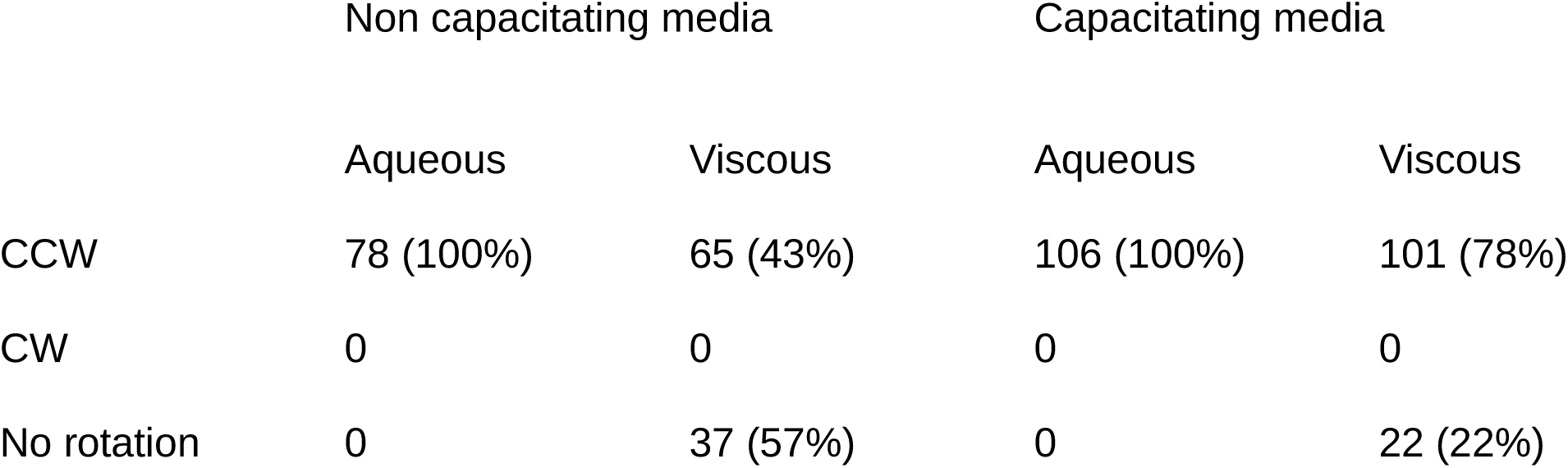
Spin direction evaluation of 409 human spermatozoa while freely swimming in 3D; 180 under non-capacitating conditions and 229 in capacitating conditions (both groups in aqueous and viscous media, see Materials and Methods). All human sperm in aqueous media (non-capacitating or capacitating) spun CCW direction when seen from head to tail regardless of the experimental condition, while in viscous and non-capacitating medium, 57% of the analyzed sperm did not spin at all, while the remaining 43% spun in the CCW direction. Note that in capacitating conditions these percentages inverted: 22% of the analyzed sperm did not spin at all, whilst 78% spun with the conserved CCW direction.

## Discussion

Sperm flagellum elastohydrodynamics, mathematical modelling and image analysis have indicated that the sperm head movement is highly dependent on the nature of forces and torques imposed by the beating flagellum and subsequent interactions with the local environment, among many other factors (Gadêlha et al., 2010; Gadêlha et al., 2019; Smith et al., 2009b; Ishimoto et al., 2017; Gaffney et al., 2011). Indeed, self-organization flagellar control models have demonstrated that mechanical attachment of the head (clamped or hinged head conditions) can even dictate the travelling wave direction of the flagellum (Riedel-Kruse et al., 2007; Camelet and Jülicher, 2020; Oriola et al., 2017; Sartori et al., 2016). These matters highlight the critical importance of directly establishing the head movement and its rotations in 3D. Furthermore, despite this critical importance and recent advances in high-speed 3D imaging and 3D microscopy, for decades there is still no consensus as to what direction human sperm spin during free-swimming motion (Muschol et al., 2018). Reports include observations of: mono-directed CW or CCW (Linnet, 1979; Smith et al., 2009b; Phillips, 1983; Woolley, 1977), bi-directed (Ishijima et al., 1992; Dardikman-Yoffe et al., 2020; Drake, 1974), and even intermittently directed head spinning (Bukatin et al., 2016). Against this background, direct detection of sperm head spinning direction and its methodology are still lacking in the literature.

Phillips, 1983 has suggested that mammalian sperm spinning direction could be easily detected with 2D bright field microscopy, by exploiting the fact that the sperm head is flattened, and thus produces blinking “flashes of light” as the head spins around its swimming axis (Phillips, 1997). It has been observed that the ‘flash of light’ travels from left-to-right when the sperm head-to-tail is aligned with the vertical axis (similarly to the sperm head orientation depicted Figure 2, and thus CW head spin was inferred. This however postulates that such ‘flash of light’ *moves in the same direction as the sperm head spinning*. We have demonstrated here this to be inconsistent with direct detection of the head spinning, see Figure 7. Instead, we have found that the true head spinning direction is opposite to the observed left-to-right movement of this “flash of light”, which similarly to Phillips, 1983 also moves from left-to-right, as depicted in Figure 7. We have shown that this mismatch of movement between optical brightness and the object spinning motion is accounted for by spherical aberration effects of the lens. Furthermore, translucent objects, such as the sperm head, are equally prone to perception bistabilities Liu et al., 2012 that equally obscure the true head spinning direction with 2D microscopy, in addition to other unknown image inversions within microscopy systems (see Methods). These uncertainties, together with the combined use of 2D views of the sperm’s head trajectory as a proxy to derive flagellar beat information, may have significantly contributed to the contradictory observations that are documented in the literature regarding the sperm head spinning direction.

In the present study, we evaluated the head spinning direction of more than 400 spermatozoa. Sperm suspended under four conditions were tested: in non-capacitating and capacitating solutions of normal and high viscosity. The high viscosity value was chosen to emulate that of the female cervical mucus (see Methods and Suarez, 2016) and sperm were incubated in capacitating media for 6 hours. One hundred percent of the spinning spermatozoa, in all experimental conditions, spun CCW, as seen from head-to-tail. It is important to mention that the spinning direction of each single spermatozoon was evaluated for a mean time of 3.4 s. A longer temporal analysis would be desirable to evaluate whether spinning direction changes over periods longer than 3 sec, though unfortunately not possible with our present high-resolution 4D set-up. Taken altogether, the analyzed time of the 409 free-swimming sperm totalled 23 minutes, with no directional change observed. Human spermatozoa in aqueous media, independently of their capacitation state, all spun CCW. In contrast, in high viscosity, 57 % of spermatozoa in non-capacitating media did not spin at all, and interestingly most of them recovered their CCW rotation (80 %) when incubated in capacitating media. This striking head spinning recovery phenomenon, as well as how sperm head spinning motion and flagellar rolling influence swimming trajectories in 3D remain to be fully explored.

## Conclusions

We studied here a half-century old problem by exploiting unique switch in contrast due to spherical aberration effect in brightfield microscopy. We have shown that 2D microscopy alone cannot distinguish spinning direction in axis-symmetric, streamlined, translucid human swimming sperm; as well, methods employing such imaging technique may need reassessment, whilst no methodology are currently available to directly measure human head spinning. Indeed, previous studies mostly used visual inspection from video-microscopy images or indirect measurement using flagellar tracing in 3D. We showed that contrast inversion can be exploited to track head spinning but this requires finding the appropriate focal plane in which the sperm head is, centred exactly at the focal plane; a challenging task for freely-swimming spermatozoa as the head moves up and down during cell progression. This is resolved by using a multi-plane detection system. The methodology was validated as coherence and consistence prevailed between optics theory and direct tracking of different sperm cells with particles attached to neck and head. We have observed that human sperm head spins with a robust, conserved, and recoverable counterclockwise spinning direction (when viewed from head to tail). Our observations reconcile structural information of mammalian sperm that observe flagellar architecture with conserved chirality in the axonemal driving unit with a conserved direction of spinning for human sperm, regardless of viscosity and capacitating conditions. The ability of human sperm to fertilize may be intimately related with head spinning, as capacitating medium were observed to excite a larger proportion of the population to spin. At last, the proposed methods can be applied to other free spinning objects and microorganisms that possess similar axis-symmetric body architecture to human spermatozoa.

## Materials and Methods

### Ethical approval for the human semen samples

The bioethics committee of the Institute of Biotechnology, UNAM approved the proposed protocols for the handling of human semen samples. Donors were properly informed regarding the experiments to be performed and each donor signed and agreed to a consent form. All samples fulfilled World Health Organization requirements for normal fertile semen samples.

### Media

HTF (human tubal fluid) medium was used in this study. Non-capacitating HTF (pH 7.4) contained (mM): 4.7 KCl, 0.3 KH_2_PO_4_, 90.7 NaCl, 1.2 MgSO_4_, 2.8 Glucose, 1.6 CaCl_2_, 3.4 sodium pyruvate, 60 sodium lactate and 23.8 HEPES. Capacitating medium (pH 7.4) was HTF medium supplemented with 5 mg/ml BSA and 2 mg/ml NaHCO_3_. Capacitating recording medium (pH 7.4) was only supplemented with 2 mg/ml NaHCO_3_, BSA was not added. For viscous medium, 1% methyl cellulose was added to the non-capacitating or capacitating recording medium, depending on the experimental condition.

### Biological preparations

Semen samples were obtained by masturbation from healthy donors after 48 h of sexual abstinence. Highly motile sperm were recovered after the swim-up protocol. Briefly, 300 µl of semen were placed in a test tube, then 1 ml of non-capacitating or capacitating medium, depending on the experimental condition, was added on top. Tubes were incubated at a 45° angle, at 37°C in a humidified atmosphere of 5% CO_2_ and 95% air during 1 h. Then, sperm from the medium on top were collected and concentration was adjusted to 10^6^ cells/ml. To promote *in vitro* capacitation, sperm in capacitating medium were incubated for an additional 5 h. Recordings were performed in (a) non-capacitating aqueous and viscous medium, and (b) capacitating aqueous or viscous medium.

### Sperm samples

A total of 409 freely swimming spermatozoa (30 samples -one per day-from 9 different donors) were analyzed: (a) 180 non-capacitated (78 in aqueous media and 102 in viscous media) and (b) 229 in capacitating media (106 in aqueous media and 123 in viscous media).

## 3D Imaging Microscopy

Multifocal plane stacks were acquired with the system originally described in Corkidi et al., 2008, consisting on an inverted Olympus IX71 microscope, mounted on an optical table [TMC (GMP SA, Switzerland)], reconfigured with a piezoelectric device P-725 (Physik Instrumente, MA, USA) which periodically displaces a high magnification 100x objective (Olympus UPlanSApo 100x/1.4 na oil objective) at a frequency of 80Hz with a z displacement of 20 mm. A high-speed camera NAC Q1v (Nac Americas, Inc., USA) acquired images at a rate of 8000 fps with 640 x 480 pixels resolution. Every rising movement of the piezo device (half cycle i.e., 1/160 sec) contains 50 different focal planes (1 image per focal plane). The high-speed camera can store 28,000 images (RAM is 8 Gb) per cell, thus recording spinning motion for a total of 3.5 second s. Since in this work the rotation of the sperm head is a critical aspect, every possible inversion in each single element conforming the optical-electronical pipeline’s path had to be carefully considered (inverted microscope, camera driver setup, image processing software and for visualization -Fiji, Matlab, Paraview, etc.-). As a control test, we have placed in the microscope stage (using a 4x objective) a known pattern (a character R in a piece of paper) viewing the front face of the objective (upside down if seen by the top of the microscope). We have verified that the character appeared upright in the computer screen and that it moved accordingly with the horizontal and vertical stage movements (seen the stage from the bottom-top direction where the objective is placed).

## Acknowledgements

The authors thank Paulina Torres for helpful assistance with experimental procedures and Shirley Ainsworth for bibliographic support. The help of Juan Manuel Hurtado, Roberto Rodríguez, Omar Arriaga and Arturo Ocádiz regarding computer services is acknowledged.

## Author contributions

Conception and design: G.C., F.M., H.G., A.D.; Acquisition of data: G.C., F.M., A-L.G.-C., Methodology: G.C., F.M., P.H.-H., A-L.G.-C, B.N.C., H.G., A.D.; Software: G.C., F.M., P.H.-H., B.N.C.; Resources: G.C., A.D.; Writing - review & editing: G.C., F.M., B.N.C., H.G., A.D.; Supervision, funding and project administration: G.C., A.D.

## Supporting Information

### Invariance of the method to sperm orientation

To investigate whether the direction of sperm rotation is not an artifact of the illumination setup of the microscope i.e., that the bright border of the sperm head edge is not an effect of the illumination setup, we analyzed the rotation of non-capacitated sperm swimming in four different directions in a Cartesian plane. As we explained previously, in the sequence shown in Figure 6, it is clearly seen that the border of the narrowest part of the head is naturally marked with a bright composed semi-circle. This bright feature turns in the direction of the sperm head (note that this bright border is located behind the sperm head, as explained before). We have verified that independently of the swimming trajectory of sperm, the evolution on time of this bright semi-circle clearly defines the rotating direction of the sperm head. Supplementary Figure S1 shows two consecutive time-points of 4 different sperm with outgoing trajectories from the center of each of four Cartesian planes. As can be seen in this figure, the bright region (due to the optical spherical aberration contrast inversion effect) in each sperm always appears first on the *b* side of the head and then it moves to the opposite *b’* side at the subsequent time-point (see Figure 9). This indicates that all the sperm rotate in a CCW (since the bright border is behind the head of the sperm) direction independently of the direction of the sperm trajectory (four quadrants).

**Figure S1.**
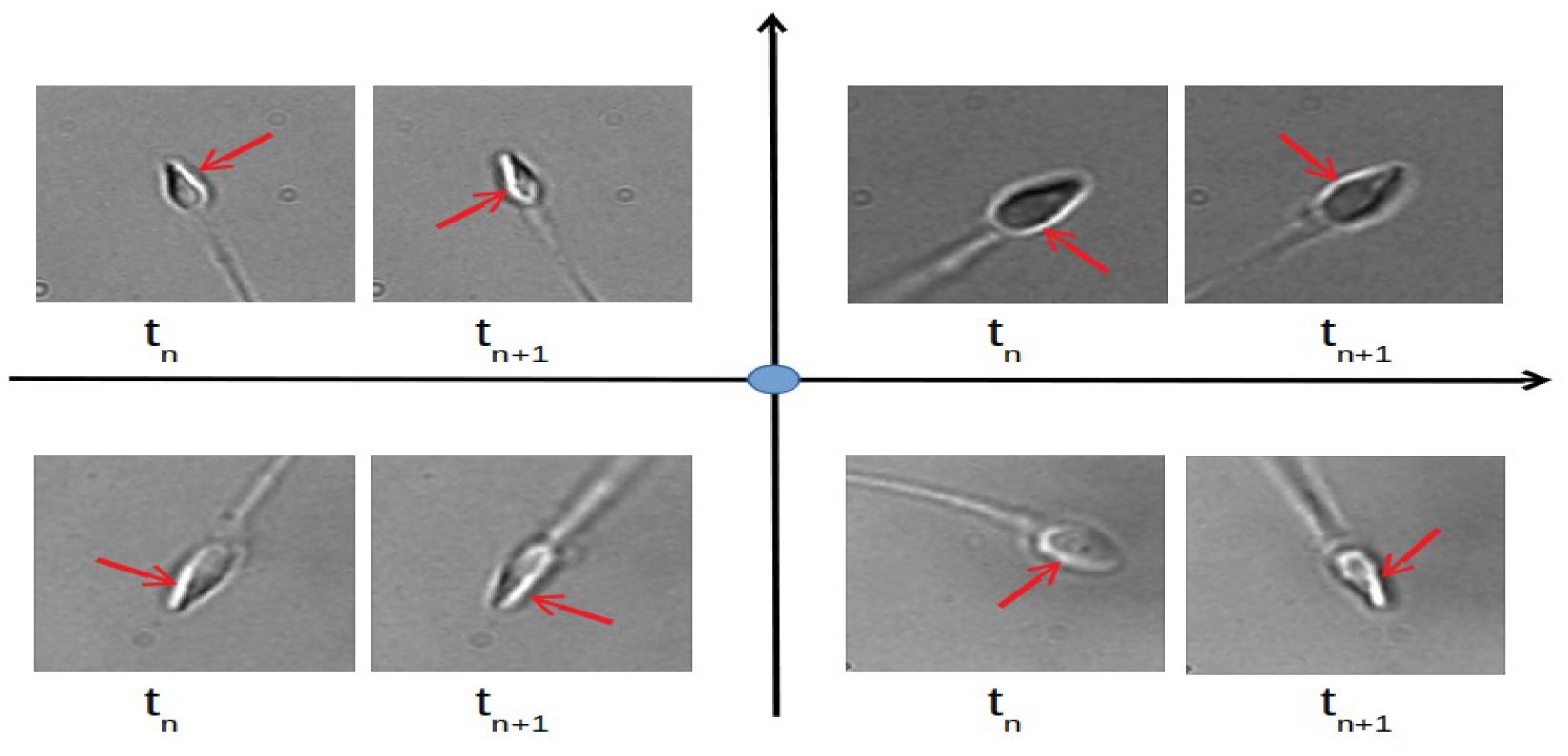
Four different non-capacitated sperm with outgoing trajectories from the center of each of the four Cartesian planes (blue circle). Two different time-points (t_n_ and t_n+1_) are shown for each sperm. The bright region (due to the optical spherical aberration contrast inversion effect) always appears first in the left side of the head and then moves to the right side at the subsequent time-point (from head to tail) indicating a CCW head spinning as shown in Results. The red arrows indicate the bright border in each image.

